# Matrisome gene expression is altered during non-allometric heart growth in genetically enlarged *Drosophila*

**DOI:** 10.1101/2024.03.25.586620

**Authors:** Rachel M. Andrews, J. Roger Jacobs

## Abstract

The cardiac extracellular matrix (ECM) is critical to defining the biophysical properties of the heart that adapt to changing stresses with growth or disease. The ECM is commonly dysregulated in chronic disease such as hypertension, diabetes, and cardiomyopathies, often leading to the development of fibrosis. There are no treatment options to address most ECM cardiomyopathies, but developing therapeutic targets necessitates an understanding of the regulation of ECM remodelling. Here, we employ a larval *Drosophila* overgrowth model (“giant larvae”) to overload the heart and alter ECM remodelling *in vivo*. These larvae grow to immense sizes without exhibiting hallmarks of obesity. Remarkably, cardiac ECM organization scales allometrically despite overload. The main effect observed is a change in Collagen fibril thickness, possibly reflecting changes to tension in the system. Overgrowth-induced changes in gene expression similarly suggest changes in Collagen assembly, such as a dramatic increase in LOXL2, the main Collagen crosslinking enzyme. This could indicate that larvae may compensate for the stress of overgrowth by stabilizing the Collagen network. The enlarged hearts of giant larvae cannot contract fully at systole. Taken together, this reveals non-allometric changes to cardiac form and output with increasing body size. Overall, our overgrowth model presents an intriguing opportunity to examine the ability of a system to tolerate overgrowth without the metabolic inputs of obesity.

## Introduction

Cardiovascular disease (CVD) is the leading cause of death worldwide, and rates are increasing globally (World Health Organization 2020). The most common risk factors for the development of CVD include aging, obesity, and diabetes. However, there are a wide variety of conditions that are at increased risk of developing CVD, including those with chronic kidney disease, inflammatory bowel disease, Marfan syndrome, and acromegaly. CVD is the leading cause of death for all of these conditions (Jankowski et al. 2021; Follin-Arbelet et al. 2023; Schicho, Marsche, and Storr 2015; Vanem et al. 2018; Kamenický, Maione, and Chanson 2021). In some conditions, like acromegaly, even individuals with pharmacologically managed disease remain at increased risk of developing CVD (Wolters et al. 2020). This highlights the importance of studying the development and progression of cardiac dysfunction and CVD.

One of the often-overlooked components of CVD is the contribution of the cardiac extracellular matrix (ECM). In a healthy system, the ECM acts as a protein scaffold that supports tissue function (Li, Zhao, and Kong 2018). The ECM is predominantly made up of Collagens, as well as proteoglycans and glycoproteins that form a highly organized network around the tissue (Jourdan-LeSaux, Zhang, and Lindsey 2010). These proteins adhere to one another by covalent crosslinks that may be formed either enzymatically or nonenzymatically (Cox and Erler 2011; Pehrsson et al. 2021). One of the main contributions of the ECM to the tissue is the regulation of biophysical properties like elasticity. This is of special importance in the heart, which must be elastic enough to contract continuously in order to maintain output, yet strong enough to support heart shape. In disease states however, dysregulation of the ECM leads to increased deposition of matrix proteins, particularly Collagens, which causes an increase in the stiffness of the heart itself (Cox and Erler 2011; Hughes and Jacobs 2017). Over time this can have severe functional consequences, as Collagen is noncontractile and can also disrupt cell-cell connections and nerve impulses through the heart (Travers et al. 2016). Pathological increase in matrix deposition is called fibrosis and is considered a hallmark of CVD (Meschiari et al. 2017; Travers et al. 2016). An additional concern is the progressive nature of fibrosis. The presence of fibrotic deposits is enough to cause the formation of more fibrotic tissue, so over time stiffness compounds to further compromise function (Bonnans, Chou, and Werb 2014; Pehrsson et al. 2021). Despite the clinical importance of fibrosis to the progression of CVD, it has no treatments that have been proven effective at preventing fibrotic remodelling (Leask 2010; Travers et al. 2022).

Developing treatments for fibrosis necessitates an understanding of how the regulation of the ECM changes in disease states. Mammalian models have been limited in their ability to address these fundamental questions due to the complexity of the ECM (Diop and Bodmer 2012). The human matrisome comprises 4% of the total proteome, and any given ECM may be composed of 100-200 different proteins (Naba et al. 2012). The use of small animal models is therefore an attractive alternative for addressing broad concepts. The genetic model system *Drosophila melanogaster* has emerged in recent years for its utility in studying the cardiac ECM. *Drosophila* is the simplest model system that possesses a heart. The formation of its simple, tubular heart follows the same developmental pathways that govern formation of the human heart, and it has a low degree of genetic redundancy, making it possible to manipulate whole gene families (Diop and Bodmer 2012; Hughes and Jacobs 2017). It has previously been reported that arresting *Drosophila* larvae in the growth phase generates larvae that grow indefinitely and reach an immense body size (Zeng et al. 2020). This model presents an intriguing opportunity to examine how the heart and the cardiac ECM adapts in the face of cardiac overload, which is known to modify ECM remodelling (Frangogiannis 2017; Hutchinson, Stewart, and Lucchesi 2010).

Here, we report that the *Drosophila* overgrowth model (referred to hereafter as “giant larvae” or “giants”) does not exhibit hallmarks characteristic of obesity. Giant larvae are significantly larger than their control counterparts, but do not have elevated triglyceride levels or increased lipid droplet size. These larvae also possess a remarkably conserved spatial organization of the cardiac ECM protein Pericardin, a heart specific Collagen found in *Drosophila*. The heart is able to adapt to this increase in body size without visible defects in the matrix. However, the heart itself is enlarged and demonstrates an inability to contract fully. Overall, cardiac morphology is preserved but cardiac output does not scale allometrically. Gene expression analysis revealed significant upregulation of the matrix crosslinking enzyme LOXL2, suggesting that giants may compensate for an increase in body size by altering the biophysical properties of their ECM. As with fibrosis, the overgrowth response reflects increased ECM deposition and increased cross-linking, but without the pathological accumulation of inelastic scars in the matrix.

## Results

### Giant larvae do not possess characteristics of obesity

In order to determine if giant larvae are a model for overgrowth or obesity we first examined some key phenotypes associated with obesity. Overall, larvae are larger in length, width, and mass compared to a wandering third instar (**Figure 1A, Figure S1**). Giant larvae are also less active than parental controls, and never initiate pre-pupation wandering behaviour. This was also observed previously (Zeng et al. 2020). Larval mass was significantly elevated, with giant larvae attaining a mass over twice that of controls (**Figure 1B**). Triglyceride (TG) levels however were significantly lower in both male and female giant larvae (**Figure 1C)**. Elevated TG levels have previously been noted in response to high fat diet (HFD) treatments that result in obesity phenotypes (Birse et al. 2010; Guida et al. 2019; Andrews et al. 2023). We investigated further by labelling lipid droplets (**Figure 1D**) within the fat body and quantifying their diameter. We find that lipid droplet diameter is reduced in females and to a lesser extent in male giant larvae (**Figure 1E**). Taken together, these results suggest that this is a model for overgrowth rather than obesity, with the caveat that some changes may reflect the delay in tissue metamorphosis.

**Figure 1:**
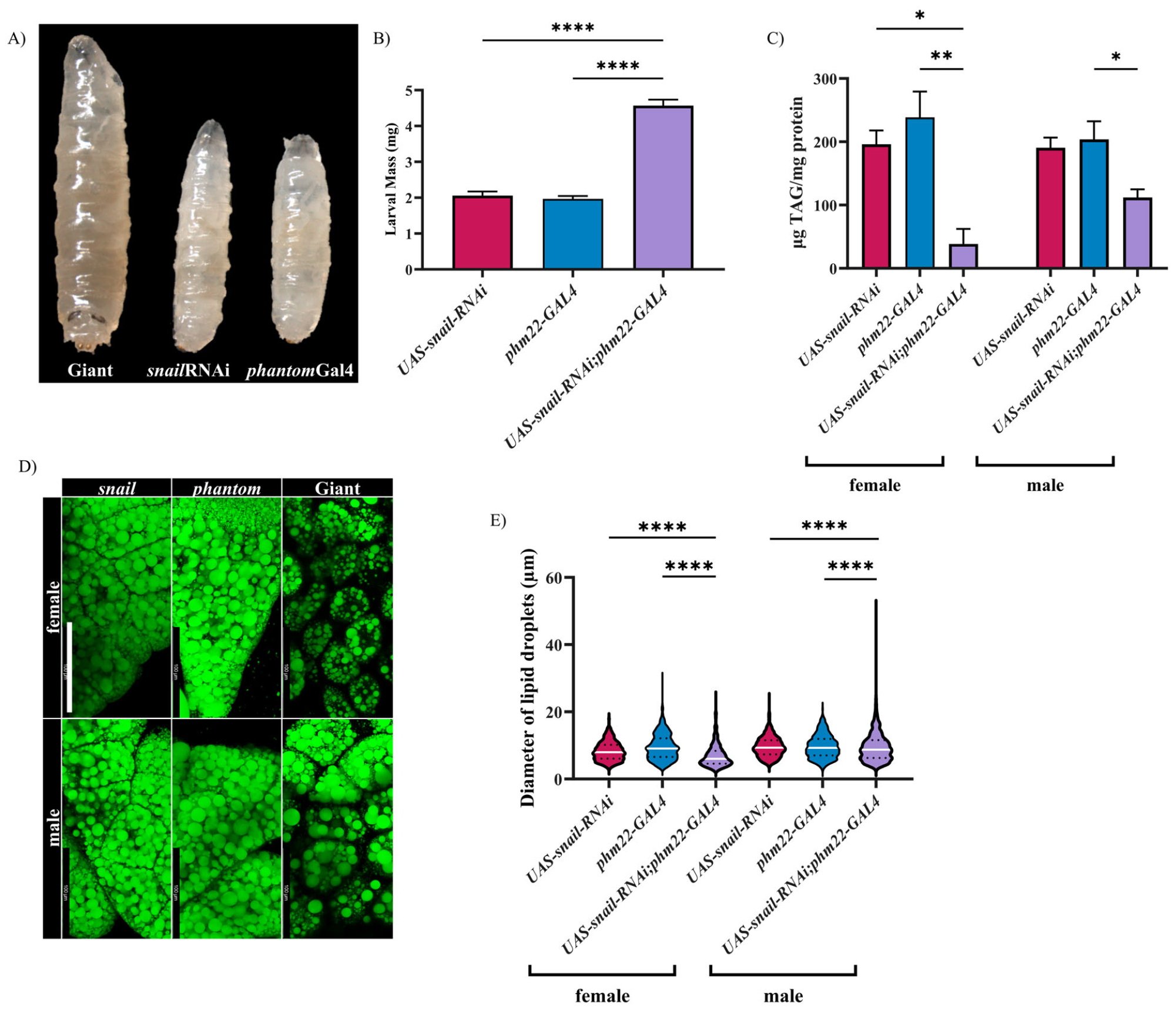
Giant larvae attain large sizes but do not demonstrate hallmarks of obesity. Giant larvae are substantially larger than parental controls at late third instar (A). Mass of giant larvae is significantly elevated compared to parental controls (B), but they have lower triglyceride levels (C). Fat body morphology is abnormal in giant larvae, with smaller lipid droplets contained compartments small sections of the fat body (D,E). Error bars in B and C are SEM. White lines in E represents the median, dotted lines represent quartiles. *=p<0.05, **=p<0.01, ***=p<0.001, ****=p<0.0001 (n>15 for larval mass, n=3 for TAG, n>5 fat bodies imaged)

### Giant larvae scale the cardiac ECM remarkably well despite overgrowth

We next aimed to determine how the cardiac ECM of the giant larvae adapts to the increase in body size. We examined the cardiac ECM by immunolabelling of Pericardin, a Collagen-IV like protein that is specific to the heart in *Drosophila* (Wilmes et al. 2018). The Pericardin network showed a remarkable tolerance for this overgrowth condition (**Figure 2)**. In controls, this network has a honeycomb appearance, with fibres pulled away laterally from the heart by the alary muscles (**Figure 2A-B’**). The appearance of the matrix was similar in giants (**Figure 2C-C’**). We previously reported defects in matrix organization with high fat diet treatments (Andrews et al. 2023), so we sought to determine if there were any organizational defects detectable in our overgrowth model. To quantify the organization of the Pericardin fibrils we utilized the Fiji macro TWOMBLI to determine the degree of alignment of fibres within the matrix. Analysis of the Pericardin fibres (see methods) revealed no significant changes in alignment between giants and controls (**Figure 3A)**. However, giants do possess thinner Pericardin fibrils than controls (**Figure 3B)**. The distribution of fibril thickness in both female and male giants was skewed towards smaller fibre widths (**Figure 3C, D**). Fibre thickness was comparable between the two parental controls. This alteration in giants could reflect increased ECM tension due to a larger body size, thus stretching fibres to cover more area. Overall, it appears that heart morphology and ECM organisation adapts remarkably well to increasing body size.

**Figure 2:**
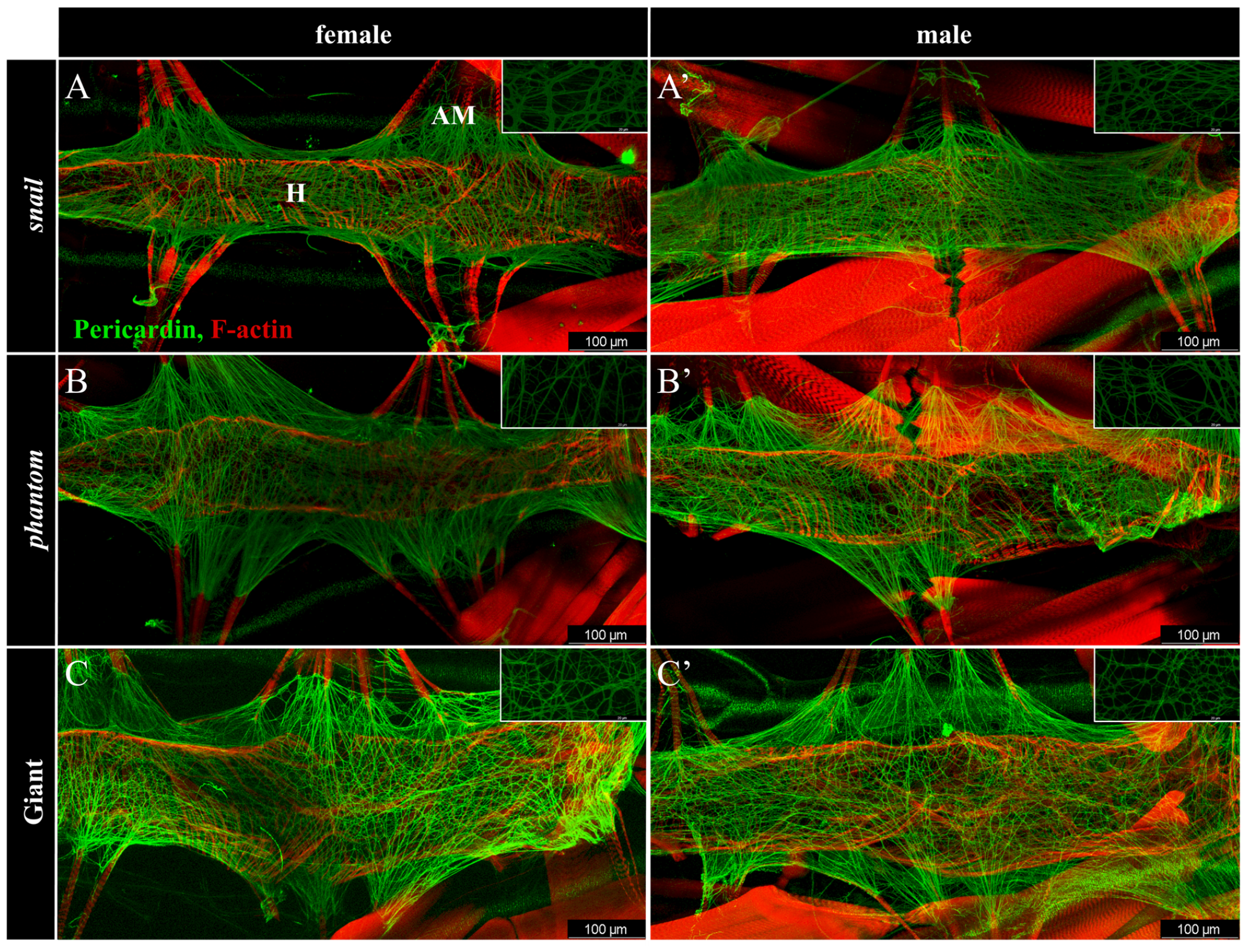
The organization of the cardiac ECM of giant larvae is remarkably conserved. Parental controls (A&A’, B&B’) show the normal Pericardin network, with a honey comb organization and the matrix being pulled away from the heart tube and anchored in the alary muscles. Giant larvae (C&C’) reveal plasticity in the organization of Pericardin, with thinner fibres (see inset). Pericardin in green, F-actin labels muscles in red. All images are oriented with anterior to the left, posterior to the right. In panel A, H labels the heart tube, AM an alary muscle. Insets 63x with 4x zoom. Scale in A is 100µm, scale in A inset 20µm. n>10 for all groups.

**Figure 3:**
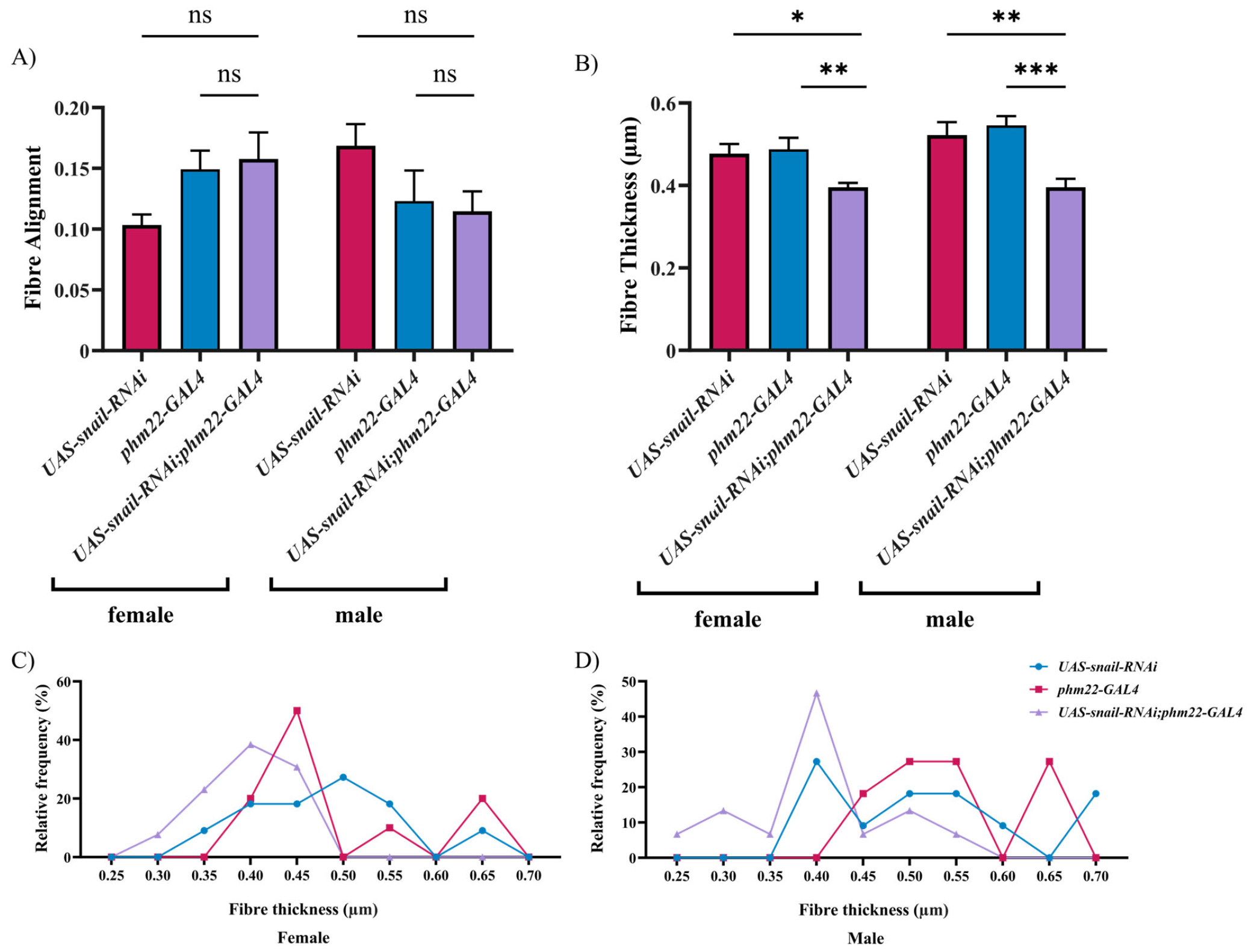
Pericardin matrix is not aligned differently in giant larvae but fibrils are thinner. Fibre alignment scores were unchanged between controls and giants (A), but the thickness of Pericardin fibres was reduced in giants (B). The distribution of fibre thicknesses was similar in control genotypes and skewed toward thinner fibres in giants (C, D). Error bars in A and B are SEM. *=p<0.05, **=p<0.01, ***=p<0.001 n>10 individuals for all groups.

### Cardiac function does not scale with increased body size

Live imaging of giant larval hearts was conducted using optical coherence tomography (OCT) (**Figure 4A**). Giant larvae were found to possess increased systolic and diastolic areas (**Figure 4C**), corresponding to an increase in stroke volume (**Figure 4D)**. Stroke volume was 1.79 fold increased in females, and 1.77 in males. However, the observed increase in mass of giant larvae is 2.22 (n=25, from Figure 1B), suggesting that the increase in stroke volume may not be enough to compensate for increased body size. Additionally, the percentage of the cross-sectional area of the body cavity that the heart occupies at diastole is significantly elevated in giant larvae (**Figure 4B**). This suggests that the heart is growing hyperallometrically with increasing body size, but proportional output is not maintained, perhaps due to an inability to contract the heart as effectively at systole.

**Figure 4:**
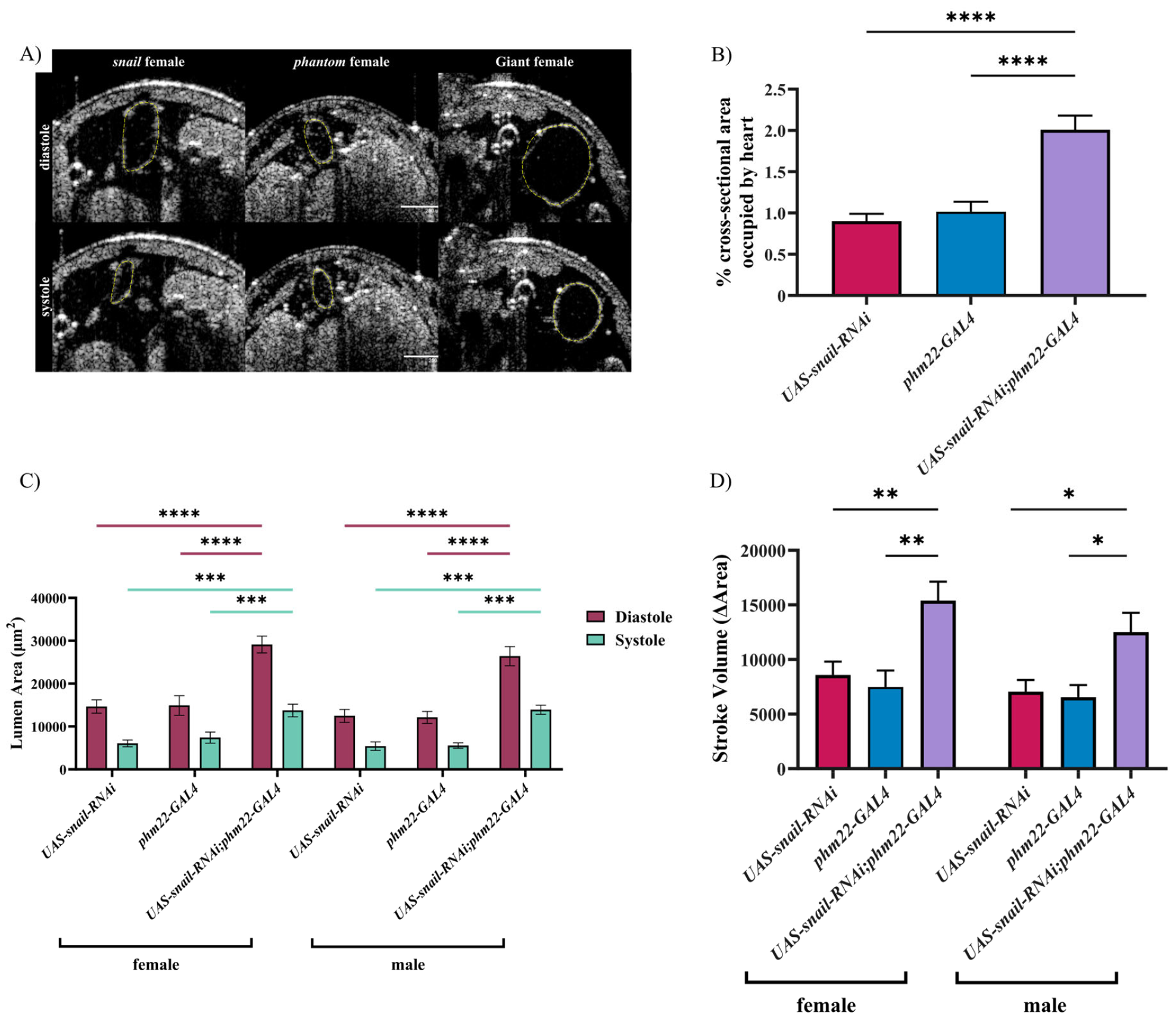
Functional analysis of larval hearts reveals disproportionately enlarged hearts in giant larvae. Optical coherence tomography (OCT) imaging reveals enlarged diastolic and systolic volume in giant larvae (outlined in yellow in A). The percentage of the body cavity that the heart occupies at diastole is also significantly elevated (B). Both female and male giant larvae have significantly increased diastolic and systolic volume (C). This corresponded to an increase in stroke volume (D), but this increase is smaller than the magnitude of the increase in body size. Error bars in B, C, and D are SEM. *=p<0.05, **=p<0.01, ***=p<0.001, ****=p<0.0001 n>10 for all groups.

### Gene expression in giant larvae is different from high fat diet treatments

Organization of the cardiac ECM was preserved in giant larvae but heart growth did not increase proportionally with body size. We therefore sought to determine what mechanism of accommodation to overgrowth could be revealed by the expression levels of genes related to ECM assembly. We also quantified expression of some genes involved in fat metabolism to determine if giants are metabolically affected by their increased body size. We compared giant larvae to wandering third instar parental controls, as well as to *y*^*1*^*w*^*1118*^ high fat diet (HFD) treated wandering third instars (Andrews et al. 2023). The parental controls revealed some variation in expression of core ECM proteins, including Pericardin, and both Collagen-IV subunits (*viking* and *Cg25c* or *Col4a1*). Giant larvae tended to follow the expression pattern of one parent or the other, with the exception of giant males showing increased Pericardin expression (**Figure 5A, C**). ECM network cross-linkers Nidogen and Perlecan showed opposite expression patterns, with giant larvae upregulating *nidogen* but downregulating *perlecan*. The metalloproteinase *MMP2* and its inhibitor *TIMP* were both also elevated **(Figure 5A, C**). HFD treated individuals did not mirror these changes, exhibiting decreased *pericardin* expression, and in females increased *cg25c* (*Col4a1*) (**Figure 5B, D)**. This indicates that HFD treatments and giant larvae regulate their ECMs differently. One notable finding was an enormous increase in expression of *LOXL2* in both female and male giant larvae **(Figure 5A, C**). Lysyl oxidase (LOX) family members are the main Collagen crosslinking enzymes. Female giant larvae showed a larger increase (∼24 fold overexpression) compared to males (∼8 fold overexpression). This trend was not observed in HFD treated larvae (**Figure 5B,D**). Overall, this suggests that ECM regulation in giant larvae is affected more dramatically than in HFD treatments, with opposing expression patterns for genes that have previously been shown to contribute to similar processes, like Nidogen and Perlecan, that have both been implicated in increasing stability of the matrix (Matsubayashi 2022).

**Figure 5:**
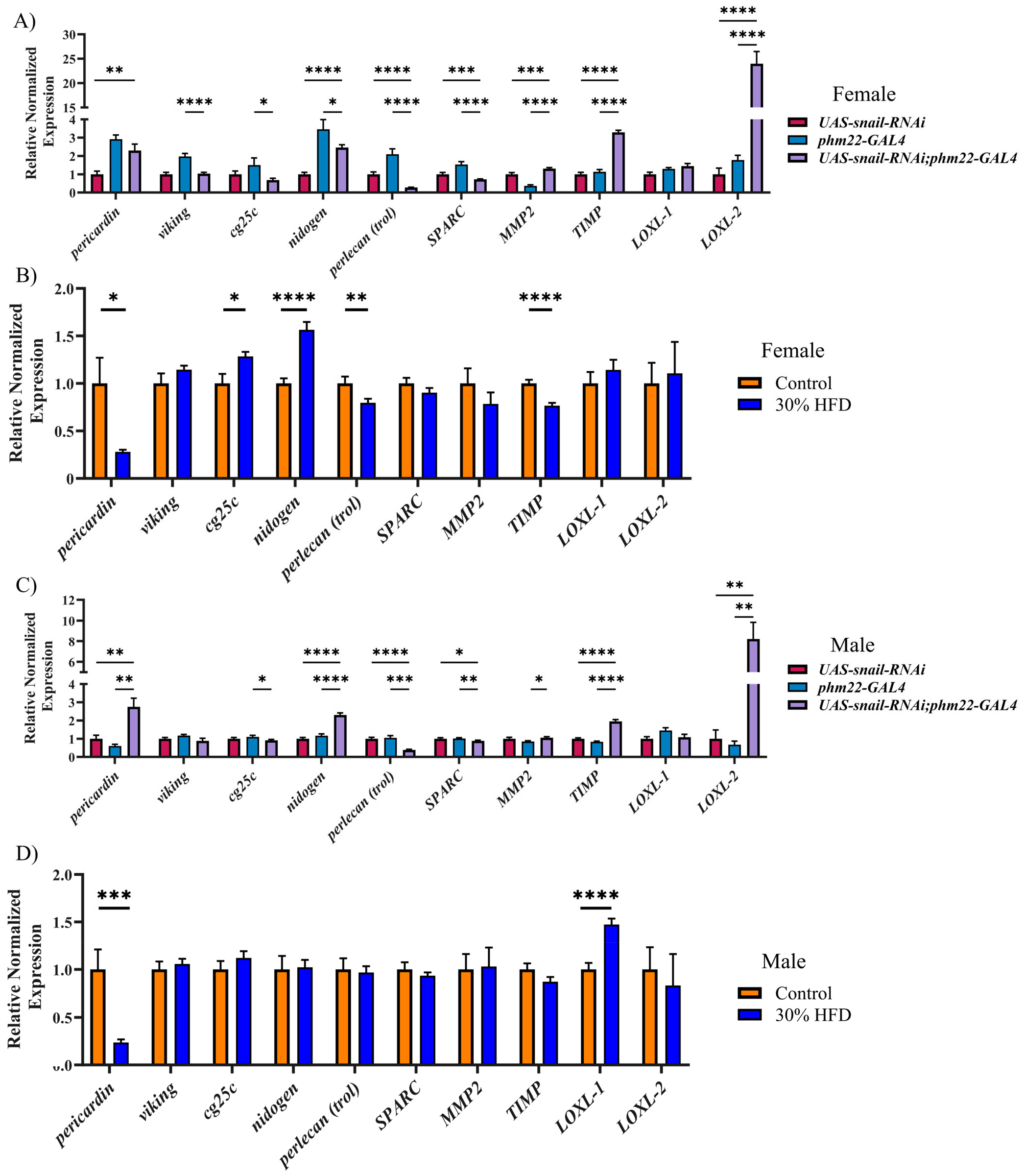
Gene expression in giant larvae differs from larvae fed a high fat diet. Giant larvae have many changes in gene expression of ECM components, and regulators (A,C). Female giant larvae exhibit an enormous increase in expression of the crosslinking enzyme LOXL2 (A). Males have a similar increase and also show increased Pericardin gene expression (B). This is in contrast to the high fat diet treatment where a decrease in Pericardin expression is observed in both females and males (C, D). Other ECM components are relatively unaffected in HFD treatments (B, D). Levels of LOXL2 are unchanged in HFD treatments. Error bars are SEM. *=p<0.05, **=p<0.01, ***=p<0.001, ****=p<0.0001, n=3 biological replicates.

Gene expression of lipid metabolism genes *lsd-2, pummelig, seipin*, and *CG5966* revealed differences in between expression in giant larvae and HFD treatments (**Figure 6**). Female and male giant larvae follow similar trends, with both sexes downregulating *lsd-2*, and upregulating *pummelig, seipin*, and *CG5966* (**Figure 6A, C**). *lsd-2* is involved in preventing the mobilization of lipid stores, suggesting that giant larvae may be utilizing energy stores (Beller et al. 2010). *pummelig* mutants experience increased lipogenesis, while *CG5966* is involved in lipid catabolism. Elevated levels of these two genes suggests that lipids are being broken down for energy and that there is decreased lipogenesis in giant larvae. HFD treatments show no differences in any of these genes in males compared to controls (**Figure 6D**), while females have mild downregulation of both *lsd-2* and *pummelig* (**Figure 6B**). *lsd-2* depletion prevents lipid storage, while *pummelig* depletion accumulates fat (Beller et al. 2010; Hehlert et al. 2019). These changes could indicate an alteration in the regulation of lipid storage in HFD females compared to males.

**Figure 6:**
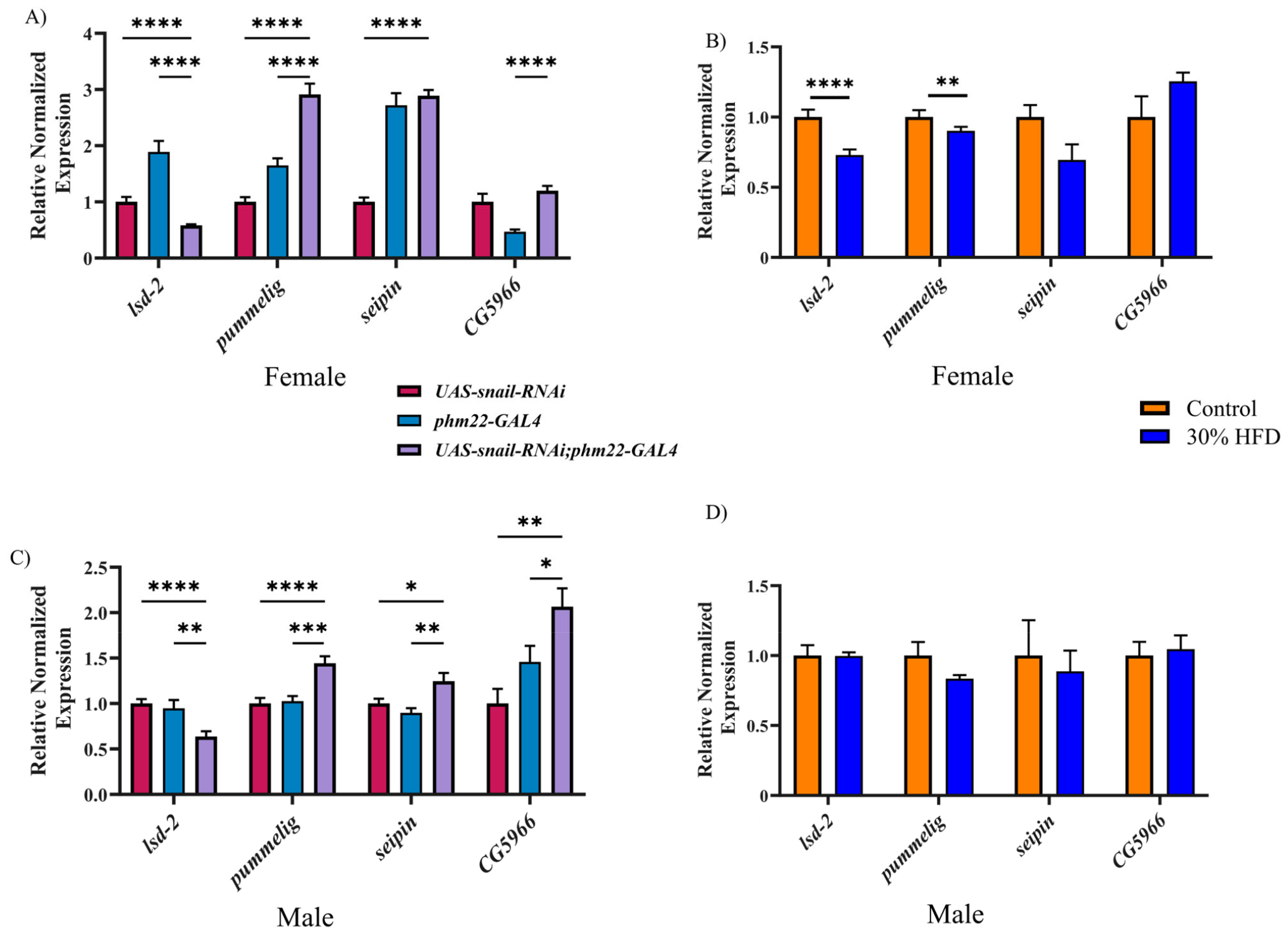
Giant larvae have significantly altered expression of genes involved in lipid metabolism. Giant larvae downregulate *lsd-2*, but upregulate *pummelig, seipin*, and *CG5966* (A, C). These changes in lipid metabolism are not observed in HFD treatment groups, with only small changes observed in females (B). Male HFD treatments show no differences in these genes (D). Female and male giant larvae follow similar trends. Error bars are SEM. *=p<0.05, **=p<0.01, ***=p<0.001, ****=p<0.0001, n=3 biological replicates.

## Discussion

Arresting *Drosophila* larvae during the growth phase of development results in larvae that are over twice the mass of a wandering third instar. Characterization of this overgrowth phenotype revealed that giant larvae grow to immense sizes without exhibiting hallmarks of obesity, including low triglyceride levels, reduced lipid droplet size, and gene expression reflecting an increase in mobilization of stored lipids, as well as decreased lipogenesis (Beller et al. 2010; Hehlert et al. 2019). This is in contrast to previous studies that have demonstrated obesity symptoms in response to high fat diet treatments (Birse et al. 2010; Guida et al. 2019; Andrews et al. 2023). This indicates that giant larvae attain their size by overgrowth, not fat storage. This presents an opportunity to examine how body size can affect the heart and other organ systems in the absence of metabolic effects, which are typically present in obesity. Additionally, acromegaly, a condition characterized by excessive growth in humans, is known to trigger cardiovascular disease (Sharma et al. 2017; Wolters et al. 2020). Cardiovascular disease is the leading cause of death in patients with acromegaly, even if their disease is medically managed (Wolters et al. 2020). Therefore having a genetic model system in which to examine adaptations made by the heart to increased body size will be informative for humans with overgrowth conditions.

Amazingly, the morphology of the cardiac ECM was largely unaffected by the overgrowth of giant larvae, despite supporting a body over twice the size. Pericardin fibre thickness was reduced in both female and male giants, with transcript abundance increased in males. Pericardin is the only fibrous, and likely the least elastic component of the *Drosophila* cardiac ECM, responsible for surrounding the heart and linking to the alary muscles that suspend it in the body cavity. Rather than growing longer fibres, overgrowth stretches those linkages and transmits the tension to the heart itself. *In vivo* imaging of active hearts revealed some physiological consequences of compensating for increased size and increased cardiac output. The heart was found to be disproportionately enlarged relative to body size with both diastolic and systolic dimensions increased. Systole may be reduced due to tension from the Pericardin network. The heart itself also occupied a larger percentage of the cross-sectional area of the body at diastole than controls. It is known in mammals that the mass of the heart scales linearly with increasing body size (Lindstedt and Scaeffer 2001). This suggests that giant larvae are experiencing hyperallometric scaling of the heart during overgrowth. The increase in stroke volume did not scale in proportion to body size. A deficit in output with a disproportionately enlarged heart provides evidence that matrix tension is increased, preventing the heart from contracting effectively at systole. Overall, the morphology of the heart adapted to overgrowth remarkably well, but the observed changes were hyperallometric and could suggest that a physiological limit has been reached.

Gene expression of important ECM components as well as ECM regulators reveals contrasting expression patterns of genes that play a role in similar processes. For example, Nidogen and Perlecan are both known to play a role in stabilizing the basement membrane (Grigorian et al. 2013; Sasse et al. 2008; Wolfstetter et al. 2019; Dai et al. 2018), but Nidogen expression was upregulated and Perlecan was downregulated during overgrowth. It has been shown previously that both of these proteins have a highly tissue specific role, and that Perlecan tends to be anchored more stably in the matrix while Nidogen remains more mobile (Matsubayashi 2022; Teuscher et al. 2024). Previous studies have shown that Collagen IV matrix turnover increases in Nidogen mutants, suggesting that Nidogen is important for stabilizing Collagen IV (Matsubayashi 2022). Our results suggest that Nidogen may be acting to stabilize the matrix during overgrowth in this system. In contrast, MMP2 expression was upregulated but TIMP was also upregulated. MMP2 is required for breakdown of the ECM during remodelling in the *Drosophila* heart and TIMP inhibits MMP2 and other proteases (Hughes et al. 2020). Taken together, this supports the idea that turnover of the matrix is dysregulated in this system. Overall, stark changes to matrisome gene expression are present in giant larvae compared to HFD treatments, which show comparatively minor changes in gene expression. The elevation of LOXL2 may be as a result of these changes to the matrisome. LOXL2 is responsible for Collagen crosslinking, which insolubilizes the matrix and makes it more resistant to degradation (Meschiari et al. 2017; Sivakumar et al. 2008). It is possible that elevated LOXL2 and increased crosslinking levels are acting to create a matrix that is both stiffer and therefore more likely to withstand mechanical strain. Increased resistance to degradation may also be compensating for increased expression of the ECM remodelling enzymes MMP2 and TIMP, which promote matrix turnover.

Here we have described a model for overgrowth that is free from confounding hallmarks of obesity. The *Drosophila* larva, an organism that normally grows 5 times in size from hatching to pupation, is able to scale its cardiac morphology well with increasing body size, even when taken beyond normal limits. The functional consequences are relatively minimal. This model presents a fascinating opportunity to determine how an organism can scale its physiology to accommodate overgrowth, which could provide insights into how interventions can be applied in a human context to improve health outcomes. The hormonally-triggered overgrowth condition acromegaly leads to increased risk factors for several diseases, even when the hormone imbalance responsible is being kept under control by treatment protocols (Wolters et al. 2020). Investigating the mechanism of response to overgrowth in this model may reveal pathways by which human overgrowth occurs, as in acromegaly.

## Methods

### *Drosophila* strains and dietary treatments

UAS-*snail*-RNAi (50003) was obtained from VDRC. UAS-*Dicer2* (BDSC 24644) was obtained from Bloomington stock centre. *phm22-GAL4* (on third chromosome) was obtained from Dr. Michael B. O’Connor. The *y*^*1*^*w*^*1118*^ background was used for high fat diet treatments. UAS*-snail-*RNAi, UAS*-Dicer2* and *phm22-GAL4* were crossed to *y*^*1*^*w*^*1118*^ for use as controls and are abbreviated here as Snail and Phm.

Flies were maintained on standard lab food, consisting of 3.6L of water, 300g sucrose (0.2M), 150g yeast, 24g KNa tartrate, 3g dipotassium hydrogen orthobasic, 1.5g NaCl, 1.5g CaCl_2,_ 1.5g MgCl_2_, 1.5g ferric sulfur, and 54g of agar. Fly food is autoclaved, cooled to 55^0^C, then 22mL of 10% tegosept and 15mL of acid mix is added before dispensing. Giant larvae were allowed to grow for 14 days at 25^0^C before being sacrificed for analysis, parental controls were taken at wandering third instar. For high fat diet treatment, 30% volume was supplemented with coconut oil and flies were maintained at room temperature (Andrews et al. 2023).

### Triglyceride assay

Triglyceride levels were measured using a serum triglyceride determination kit (Sigma Aldrich, TR0100) ((Wat et al. 2020). 5 intact third instar larvae were flash frozen in liquid nitrogen and stored at -80^0^C before sample preparation. Frozen larvae were ground with a manual homogenizer in 0.1% Tween in PBS. 20µl of buffer per larva was used. Samples were heat treated at 70^0^C for 10 minutes, then centrifuged at maximum speed for 3 minutes. 10µL of each sample was loaded into a 96 well plate in triplicate. 10µL of a glycerol standard at 2.5mg/mL, 1.25mg/mL, 0.625mg/mL, 0.315mg/mL, 0.156mg/mL, and 0mg/mL were also loaded. 250µL of free glycerol reagent was added to each well, incubated at 37^0^C, and absorbance was read at 540nm. 50µL of triglyceride reagent was then added, incubated for 10 minutes at 37^0^C, and absorbance read at 540nm. The change in glycerol levels after addition of the triglyceride reagent was calculated to determine the level of stored triglycerides in the sample. A Bradford assay was then conducted on the same samples and the level of stored triglycerides was divided by the amount of protein in the sample to control for body size.

### Dissections

#### Heart

Dissections were performed by fixing larvae dorsal down to a surface using pins (Brent, Werner, and McCabe 2009). Larvae were bathed in PBS and an incision was made at the ventral midline. The cuticle was pinned back and the gut and fat bodies were removed to reveal the heart. Control dissections were performed at third instar, after the onset of wandering behaviour. Giant larvae were dissected at day 14 post laying.

#### Fat body

Above process was followed but only the gut was removed to expose the fat bodies.

### Immunohistochemistry

#### Heart

Dissections were fixed for 20 minutes without shaking at room temperature in 4% paraformaldehyde in PBS. Specimens were then washed 3×10 minutes in PBST (0.3% Triton-X-100), before blocking for 30 minutes with NGS (1:15). Primary antibodies were incubated overnight at 4^0^C with shaking. After incubation with primary 3×10 minute washes in PBST were performed before adding secondary antibodies for one hour at room temperature. Phalloidin was added at the same time as secondary antibodies. Specimens were then washed 3×10 minutes in PBST, with a final wash in PBS to remove detergent. 50% glycerol was added for at least 3 hours, then 70% glycerol overnight. The primary antibody used was mouse anti-Prc (Pericardin, EC11, DSHB, 1:30 dilution). Secondary antibodies used were Alexa Fluor 488 anti-mouse and Alexa Fluor 647 anti-mouse (1:150 dilution). Alexa Fluor 546 and 647 Phalloidin (Thermofisher Scientific) were also used (1:75 dilution).

#### Fat body

Dissections were fixed for 30 minutes at room temperature in 4% paraformaldehyde. Specimens were washed 2×5 minutes in PBST, then incubated in 493/503 BODIPY (1:1000) for 30 minutes. Specimens were then washed 2×5 minutes, placed in 70% glycerol, and immediately mounted for imaging.

### Imaging

A Leica SP5 confocal microscope was used to obtain image stacks. 1µm intervals between frames were used for heart dissections, 0.5µm intervals were used for fat bodies. Fat bodies were imaged from the surface to a depth of 30µm. Hearts were imaged from the ventral face of the cardiac ECM to the dorsal edge of the heart tube. Images were processed using Leica software (LAS AF), ImageJ, and ZEN blue.

### OCT imaging

Optical coherence tomography (OCT) was used to visualize the heart beating *in vivo* in real time in late third instar larvae. Larvae were adhered to a microscope slide dorsal side up before being placed under the OCT camera. B scans were taken in 3D acquisition mode using a Thorlabs OCT Telesto series TEL221PS system at the widest point of the heart chamber with the following parameters: X size 1257 pixels, 1.03mm, Y size 0, 400 frames, Z field of view 1.2mm. This gives a 20 second video with 20 frames per second. Image stacks were then exported as TIFs and processed in ImageJ (Abràmoff, Magalhaes, and Ram 2004). The cross-sectional area was measured at both diastole and systole. The difference between diastolic and systolic volumes was used as a proxy for stroke volume.

### qPCR

#### RNA extraction and RT-qPCR

Total RNA was extracted from larvae using TRIzol (Invitrogen, 15596026). Wandering third instar larvae or 14 day old giant larvae were flash frozen in liquid nitrogen in groups of 5 (n=3). Samples were stored at -80 or left in liquid nitrogen until ready to use. Samples were then ground in 800µL TRIzol, and incubated at room temperature (RT) for 5 minutes. 80µL chloroform was then added, samples were shaken for 15 seconds, incubated at RT for 3 minutes, and spun at 12,000RPM for 15 minutes at 4°C. After the spin, the supernatant was added to a gDNA eliminator column (Qiagen RNeasy plus kit, 74034) and spun at RT for 30 seconds at 10,000RPM. 600µL of 70% ethanol was added to the flow through, which was then transferred to the RNeasy spin column. This was spun for 15 seconds at 10,000RPM RT, flow through was discarded, 700µL of buffer RW1 was added, spun for 15 seconds at 10,000 RPM RT, flow through discarded, 500µL buffer RPE was added, and spun at 10,000RPM RT for 2 minutes. The column was then placed in a fresh tube, 40µL RNase free water was added, incubated for 10 minutes, and spun for 1 minute at 10,000RPM RT. Samples were then used to make cDNA using the Applied Biosystems High-Capacity cDNA reverse transcription kit (ThermoFisher, 4368814).

RT-qPCR was performed (in triplicate) using the Bio-Rad cycler CFX 96 and the Luna universal qPCR master mix (NEB, M3003X). Gene expression levels were normalized to housekeeping genes EF1, Rpl32, and α-tubulin. Primers can be found in supplemental materials.

### Quantification and statistics

Fibre alignment was quantified using using the Twombli plug-in in Fiji 2.14 (ImageJ2 ver2.9.0, http://imagej.net) (Abràmoff, Magalhaes, and Ram 2004; Schindelin et al. 2012). Parameters were adjusted to detect fibres of 7-25 units and minimum branch length of 15 units. Masks were compared against original single channel confocal images. (Wershof et al. 2021).

Fibre thickness was measured using 63x images with 4x zoom in ImageJ (Abràmoff, Magalhaes, and Ram 2004). All fibres within a 15×15µm ROI were measured.

Lipid droplet diameter was measured using the line tool in ZEN 3.4 (blue edition).

Statistical analysis of larval health (mass, triglyceride levels, lipid droplet size), Pericardin fibre thickness, and OCT measurements were performed using Graphpad Prism (v.9.5.1). Analysis of variance (ANOVA) with a multiple comparison’s test was performed. Graphs are plotted with SEM.

RT-qPCR results were analyzed using CFX Maestro 3.1 software (Bio-Rad, Canada; https://www.bio-rad.com/en-ca/product/cfx-maestro-software-for-cfx-real-time-pcr-instruments), which performed an ANOVA or a t-test depending on number of groups being compared.

## Supplemental methods

Primers used:

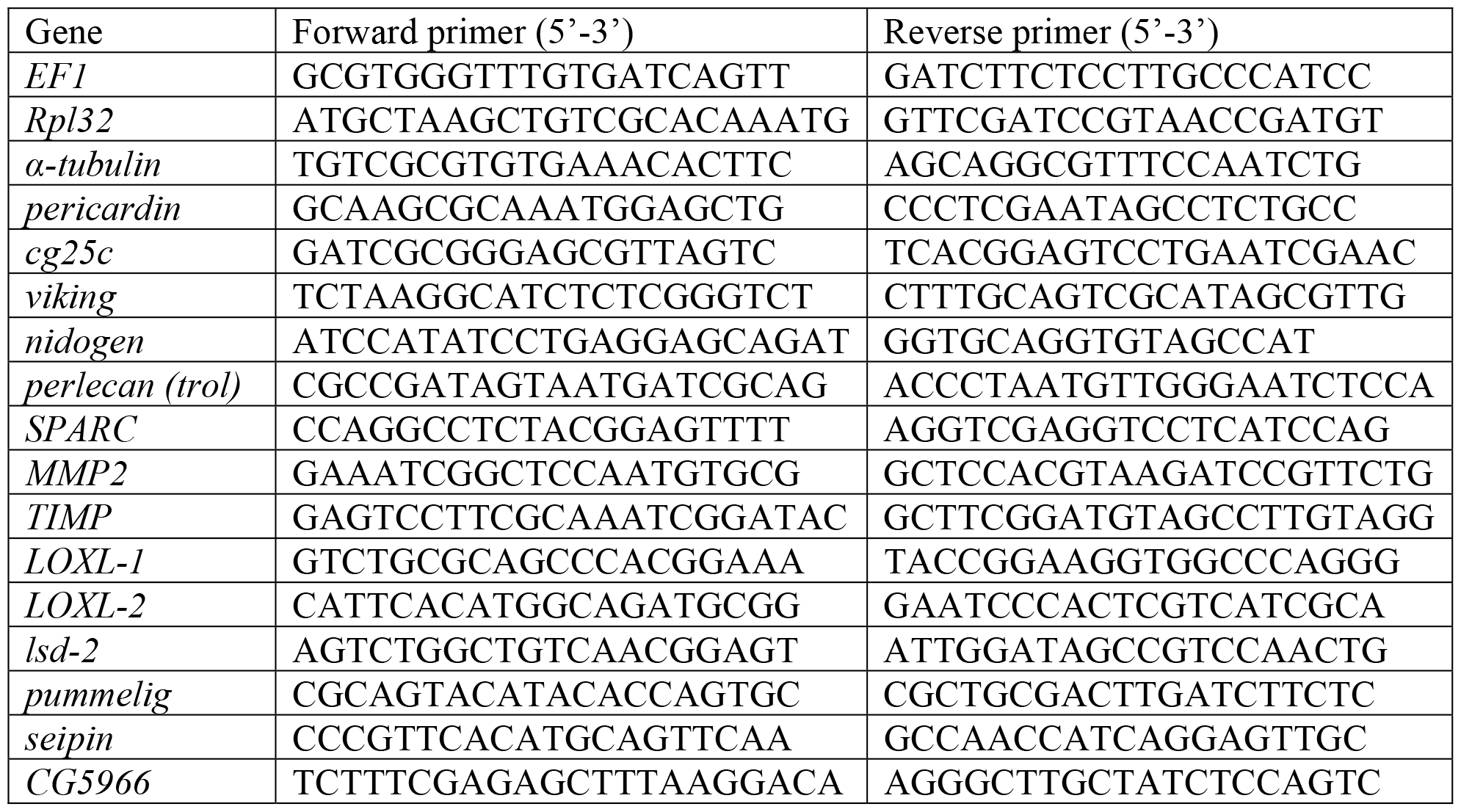

## Supplemental figures

**Figure S1:**
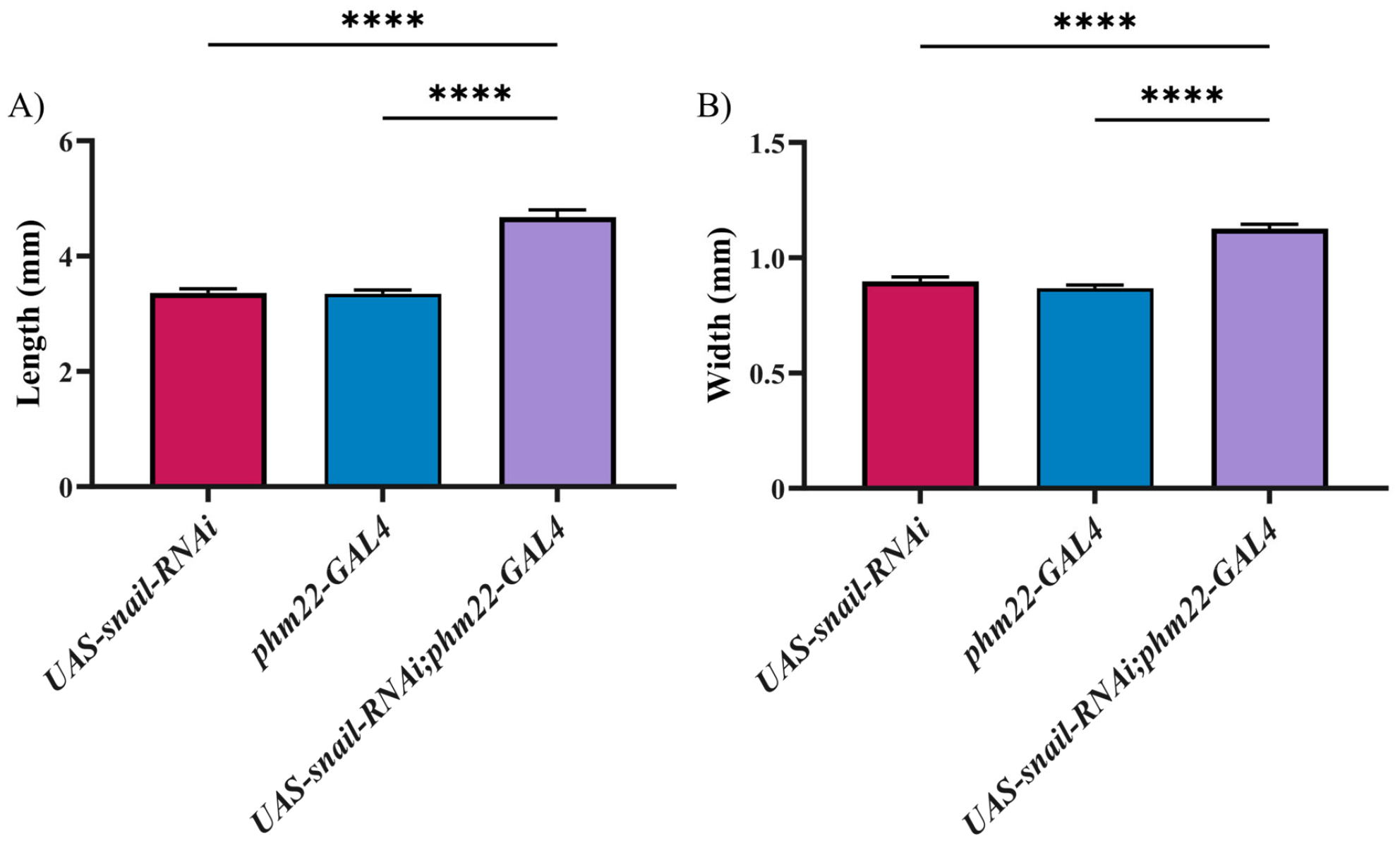
Giant larvae have significantly increased body measurements. Giant larvae have significantly increased body length (A) as well as width (B). Error bars are SEM. ****=p<0.0001

## References

Abràmoff, Dr Michael D, Paulo J Magalhaes, and Sunanda Ram. 2004. “Image Processing with ImageJ.” Biophotonics International, July.

Andrews, Rachel M., Saumya Naik, Katie Pelletier, and J. Roger Jacobs. 2023. “Cardiac Function and ECM Morphology Are Altered with High Fat Diets in Drosophila.” bioRxiv. 10.1101/2023.08.08.552539.

Beller, Mathias, Anna V. Bulankina, He-Hsuan Hsiao, Henning Urlaub, Herbert Jäckle, and Ronald P. Kühnlein. 2010. “PERILIPIN-Dependent Control of Lipid Droplet Structure and Fat Storage in Drosophila.” Cell Metabolism 12 (5): 521–32. 10.1016/j.cmet.2010.10.001.

Birse, Ryan T., Joan Choi, Kathryn Reardon, Jessica Rodriguez, Suzanne Graham, Soda Diop, Karen Ocorr, Rolf Bodmer, and Sean Oldham. 2010. “High-Fat-Diet-Induced Obesity and Heart Dysfunction Are Regulated by the TOR Pathway in Drosophila.” Cell Metabolism 12 (5): 533–44. 10.1016/j.cmet.2010.09.014.

Bonnans, Caroline, Jonathan Chou, and Zena Werb. 2014. “Remodelling the Extracellular Matrix in Development and Disease.” Nature Reviews Molecular Cell Biology 15 (12): 786–801. 10.1038/nrm3904.

Brent, Jonathan R., Kristen M. Werner, and Brian D. McCabe. 2009. “Drosophila Larval NMJ Dissection.” Journal of Visualized Experiments: JoVE, no. 24 (February): 1107. 10.3791/1107.

Cox, Thomas R., and Janine T. Erler. 2011. “Remodeling and Homeostasis of the Extracellular Matrix: Implications for Fibrotic Diseases and Cancer.” Disease Models & Mechanisms 4 (2): 165–78. 10.1242/dmm.004077.

Dai, Jianli, Beatriz Estrada, Sofie Jacobs, Besaiz J. Sánchez-Sánchez, Jia Tang, Mengqi Ma, Patricia Magadán-Corpas, José C. Pastor-Pareja, and María D. Martín-Bermudo. 2018. “Dissection of Nidogen Function in Drosophila Reveals Tissue-Specific Mechanisms of Basement Membrane Assembly.” PLoS Genetics 14 (9): e1007483. 10.1371/journal.pgen.1007483.

Diop, Soda Balla, and Rolf Bodmer. 2012. “Drosophila as a Model to Study the Genetic Mechanisms of Obesity-Associated Heart Dysfunction.” Journal of Cellular and Molecular Medicine 16 (5): 966–71. 10.1111/j.1582-4934.2012.01522.x.

Follin-Arbelet, Benoit, Milada Cvancarova Småstuen, Øistein Hovde, Lars-Petter Jelsness-Jørgensen, and Bjørn Moum. 2023. “Mortality in Patients with Inflammatory Bowel Disease: Results from 30 Years of Follow-up in a Norwegian Inception Cohort (the IBSEN Study).” Journal of Crohn’s and Colitis 17 (4): 497–503. 10.1093/ecco-jcc/jjac156.

Frangogiannis, Nikolaos G. 2017. “The Extracellular Matrix in Myocardial Injury, Repair, and Remodeling.” The Journal of Clinical Investigation 127 (5): 1600–1612. 10.1172/JCI87491.

Grigorian, Melina, Ting Liu, Utpal Banerjee, and Volker Hartenstein. 2013. “The Proteoglycan Trol Controls Proliferation and Differentiation of Blood Progenitors in the Drosophila Lymph Gland.” Developmental Biology 384 (2): 301–12. 10.1016/j.ydbio.2013.03.007.

Guida, Maria Clara, Ryan Tyge Birse, Alessandra Dall’Agnese, Paula Coutinho Toto, Soda Balla Diop, Antonello Mai, Peter D. Adams, Pier Lorenzo Puri, and Rolf Bodmer. 2019. “Intergenerational Inheritance of High Fat Diet-Induced Cardiac Lipotoxicity in Drosophila.” Nature Communications 10 (January): 193. 10.1038/s41467-018-08128-3.

Hehlert, Philip, Vinzenz Hofferek, Christoph Heier, Thomas O. Eichmann, Dietmar Riedel, Jonathan Rosenberg, Anna Takacs, et al. 2019. “The α/β-Hydrolase Domain-Containing 4- and 5-Related Phospholipase Pummelig Controls Energy Storage in Drosophila.” Journal of Lipid Research 60 (8): 1365–78. 10.1194/jlr.M092817.

Hughes, C. J. R., and J. Roger Jacobs. 2017. “Dissecting the Role of the Extracellular Matrix in Heart Disease: Lessons from the Drosophila Genetic Model.” Veterinary Sciences 4 (2): 24. 10.3390/vetsci4020024.

Hughes, C.J.R., S. Turner, R.M. Andrews, A. Vitkin, and J.R. Jacobs. 2020. “Matrix Metalloproteinases Regulate ECM Accumulation but Not Larval Heart Growth in Drosophila Melanogaster.” Journal of Molecular and Cellular Cardiology 140 (March): 42–55. 10.1016/j.yjmcc.2020.02.008.

Hutchinson, Kirk R., James A. Stewart, and Pamela A. Lucchesi. 2010. “Extracellular Matrix Remodeling During the Progression of Volume Overload-Induced Heart Failure.” Journal of Molecular and Cellular Cardiology 48 (3): 564–69. 10.1016/j.yjmcc.2009.06.001.

Jankowski, Joachim, Jürgen Floege, Danilo Fliser, Michael Böhm, and Nikolaus Marx. 2021. “Cardiovascular Disease in Chronic Kidney Disease.” Circulation 143 (11): 1157–72. 10.1161/CIRCULATIONAHA.120.050686.

Jourdan-LeSaux, Claude, Jianhua Zhang, and Merry L. Lindsey. 2010. “Extracellular Matrix Roles during Cardiac Repair.” Life Sciences 87 (13): 391–400. 10.1016/j.lfs.2010.07.010.

Kamenický, Peter, Luigi Maione, and Philippe Chanson. 2021. “Cardiovascular Complications of Acromegaly.” Annales d’Endocrinologie, 63rd International Meeting of Clinical Endocrinology - Henri-Pierre KLOTZ : Heart and Hormones, 82 (3): 206–9. 10.1016/j.ando.2020.03.010.

Leask, Andrew. 2010. “Potential Therapeutic Targets for Cardiac Fibrosis.” June 2010. 10.1161/CIRCRESAHA.110.217737.

Li, Li, Qian Zhao, and Wei Kong. 2018. “Extracellular Matrix Remodeling and Cardiac Fibrosis.” Matrix Biology 68–69 (August): 490–506. 10.1016/j.matbio.2018.01.013.

Lindstedt, S. L., and P. J. Scaeffer. 2001. “Use of Allometry in Predicting Anatomical and Physiological Parameters of Mammals.” 2001. 10.1258/0023677021911731.

Matsubayashi, Yutaka. 2022. “Dynamic Movement and Turnover of Extracellular Matrices during Tissue Development and Maintenance.” Fly 16 (1): 248–74. 10.1080/19336934.2022.2076539.

Meschiari, Cesar A., Osasere Kelvin Ero, Haihui Pan, Toren Finkel, and Merry L. Lindsey. 2017. “The Impact of Aging on Cardiac Extracellular Matrix.” GeroScience 39 (1): 7–18. 10.1007/s11357-017-9959-9.

Naba, Alexandra, Karl R. Clauser, Sebastian Hoersch, Hui Liu, Steven A. Carr, and Richard O. Hynes. 2012. “The Matrisome: In Silico Definition and In Vivo Characterization by Proteomics of Normal and Tumor Extracellular Matrices.” Molecular & Cellular Proteomics 11 (4): M111.014647. 10.1074/mcp.M111.014647.

Pehrsson, Martin, Joachim Høg Mortensen, Tina Manon-Jensen, Anne-Christine Bay-Jensen, Morten Asser Karsdal, and Michael Jonathan Davies. 2021. “Enzymatic Cross-Linking of Collagens in Organ Fibrosis – Resolution and Assessment.” Expert Review of Molecular Diagnostics 21 (10): 1049–64. 10.1080/14737159.2021.1962711.

Sasse, Philipp, Daniela Malan, Michaela Fleischmann, Wilhelm Roell, Erika Gustafsson, Toktam Bostani, Yun Fan, et al. 2008. “Perlecan Is Critical for Heart Stability.” Cardiovascular Research 80 (3): 435–44. 10.1093/cvr/cvn225.

Schicho, Rudolf, Gunther Marsche, and Martin Storr. 2015. “Cardiovascular Complications in Inflammatory Bowel Disease.” Current Drug Targets 16 (3): 181–88.

Schindelin, Johannes, Ignacio Arganda-Carreras, Erwin Frise, Verena Kaynig, Mark Longair, Tobias Peitzsch, Stephan Preibisch, et al. 2012. “Fiji: An Open-Source Platform for Biological-Image Analysis | Nature Methods.” June 28, 2012. https://www.nature.com/articles/nmeth.2019.

Sharma, Morali D., Anh V. Nguyen, Spandana Brown, and Richard J. Robbins. 2017. “Cardiovascular Disease in Acromegaly.” Methodist DeBakey Cardiovascular Journal 13 (2): 64–67. 10.14797/mdcj-13-2-64.

Sivakumar, P., Sudhiranjan Gupta, Sagartirtha Sarkar, and Subha Sen. 2008. “Upregulation of Lysyl Oxidase and MMPs during Cardiac Remodeling in Human Dilated Cardiomyopathy.” Molecular and Cellular Biochemistry 307 (1): 159–67. 10.1007/s11010-007-9595-2.

Teuscher, Alina C., Cyril Statzer, Anita Goyala, Seraina A. Domenig, Ingmar Schoen, Max Hess, Alexander M. Hofer, et al. 2024. “Longevity Interventions Modulate Mechanotransduction and Extracellular Matrix Homeostasis in C. Elegans.” Nature Communications 15 (1): 276. 10.1038/s41467-023-44409-2.

Travers, Joshua G., Fadia A. Kamal, Jeffrey Robbins, Katherine E. Yutzey, and Burns C. Blaxall. 2016. “Cardiac Fibrosis: The Fibroblast Awakens.” Circulation Research 118 (6): 1021–40. 10.1161/CIRCRESAHA.115.306565.

Travers, Joshua G., Charles A. Tharp, Marcello Rubino, and Timothy A. McKinsey. 2022. “Therapeutic Targets for Cardiac Fibrosis: From Old School to next-Gen.” The Journal of Clinical Investigation 132 (5). 10.1172/JCI148554.

Vanem, Thy Thy, Odd Ragnar Geiran, Kirsten Krohg-Sørensen, Cecilie Røe, Benedicte Paus, and Svend Rand-Hendriksen. 2018. “Survival, Causes of Death, and Cardiovascular Events in Patients with Marfan Syndrome.” Molecular Genetics & Genomic Medicine 6 (6): 1114–23. 10.1002/mgg3.489.

Wat, Lianna W., Charlotte Chao, Rachael Bartlett, Justin L. Buchanan, Jason W. Millington, Hui Ju Chih, Zahid S. Chowdhury, et al. 2020. “A Role for Triglyceride Lipase Brummer in the Regulation of Sex Differences in Drosophila Fat Storage and Breakdown.” Edited by Bassem A. Hassan. PLOS Biology 18 (1): e3000595. 10.1371/journal.pbio.3000595.

Wershof, Esther, Danielle Park, David J Barry, Robert P Jenkins, Antonio Rullan, Anna Wilkins, Karin Schlegelmilch, et al. 2021. “A FIJI Macro for Quantifying Pattern in Extracellular Matrix.” Life Science Alliance 4 (3): e202000880. 10.26508/lsa.202000880.

Wilmes, Ariane C., Nora Klinke, Barbara Rotstein, Heiko Meyer, and Achim Paululat. 2018. “Biosynthesis and Assembly of the Collagen IV-like Protein Pericardin in Drosophila Melanogaster.” Biology Open 7 (4): bio030361. 10.1242/bio.030361.

Wolfstetter, Georg, Ina Dahlitz, Kathrin Pfeifer, Uwe Töpfer, Joscha Arne Alt, Daniel Christoph Pfeifer, Reinhard Lakes-Harlan, Stefan Baumgartner, Ruth H. Palmer, and Anne Holz. 2019. “Characterization of Drosophila Nidogen/Entactin Reveals Roles in Basement Membrane Stability, Barrier Function and Nervous System Patterning.” Development 146 (2): dev168948. 10.1242/dev.168948.

Wolters, Thalijn L. C., Mihai G. Netea, Niels P. Riksen, Adrianus R. M. M. Hermus, and Romana T. Netea-Maier. 2020. “Acromegaly, Inflammation and Cardiovascular Disease: A Review.” Reviews in Endocrine & Metabolic Disorders 21 (4): 547–68. 10.1007/s11154-020-09560-x.

World Health Organization. 2020. “Global Health Estimates 2020: Deaths by Cause, Age, Sex, by Country, and by Region, 2000-2019.” Geneva.

Zeng, Jie, Nhan Huynh, Brian Phelps, and Kirst King-Jones. 2020. “Snail Synchronizes Endocycling in a TOR-Dependent Manner to Coordinate Entry and Escape from Endoreplication Pausing during the Drosophila Critical Weight Checkpoint.” Edited by Bruce Edgar. PLOS Biology 18 (2): e3000609. 10.1371/journal.pbio.3000609.

